# Leucine Aminopeptidase 3 Regulates Skeletal Muscle Mitochondrial Homeostasis with Sex-Dependent Metabolic Consequences

**DOI:** 10.64898/2026.06.20.733486

**Authors:** Shion Osana, Reiya Murakami, Ryui Natsuyama, Ayaka Tabuchi, Ryotaro Kano, Kohei Baba, Haopeng Wang, Hiroaki Takada, Naoki Suzuki, Kazutaka Murayama, Makoto Kanzaki, Yasuo Kitajima, Mizuki Sudo, Daisuke Hoshino, Ryoichi Nagatomi, Yutaka Kano

## Abstract

Skeletal muscle homeostasis depends on the coordinated regulation of protein turnover and mitochondrial quality control; however, the molecular mechanisms linking these processes remain unclear. In this study, we examined the physiological role of leucine aminopeptidase 3 (LAP3), a post-proteolytic aminopeptidase, using constitutive LAP3-deficient mice. LAP3 deficiency preferentially affected skeletal muscle, causing reduced muscle mass and mitochondrial enlargement in both sexes. Female LAP3-deficient mice also showed reduced myofiber size, impaired endurance capacity, increased energy expenditure, elevated lipid oxidation, and lipid droplet accumulation adjacent to the mitochondria. Proteomic analyses revealed remodeling of pathways related to lipid metabolism and protein homeostasis. Consistent with these findings, LAP3 deficiency increased the expression of Pink1 and Tax1bp1 and promoted the accumulation of ubiquitinated proteins, suggesting alterations in mitochondrial quality control and proteostatic regulation. In cultured myogenic cells, LAP3 localized to mitochondrial fractions, and both LAP3 knockdown and overexpression altered mitochondrial morphology. Taken together, these results identify LAP3 as a regulator of skeletal muscle homeostasis and support a role for LAP3 in linking intracellular peptide turnover to mitochondrial homeostasis, with female skeletal muscle showing greater susceptibility to LAP3 deficiency.

## Introduction

Skeletal muscle plays an important role not only in locomotor function and postural maintenance, but also in the regulation of systemic glucose and lipid metabolism^1,2^. The maintenance of skeletal muscle function depends on the preservation of myofiber integrity and mitochondrial homeostasis^3^. Disruption of these processes contributes to muscle wasting and metabolic disorders, in which impaired endurance capacity and mitochondrial dysfunction are common pathological features.

Protein turnover and myogenic differentiation are key processes underlying the maintenance of muscle integrity. The ubiquitin–proteasome system removes unwanted proteins and preserves myofiber quality^4–7^. Aminopeptidases further process proteasome-generated peptides^8,9^, which contribute to intracellular amino acid homeostasis and metabolic regulation^10–13^. Several aminopeptidases have been implicated in muscle differentiation and protein quality control^14–17^. Of these, leucine aminopeptidase 3 (LAP3) has emerged as a regulator of myogenesis. Functional studies have demonstrated that LAP3 contributes to myoblast differentiation, which supports a role for LAP3 in skeletal muscle development and maturation^18,19^.

In addition to its role in peptide processing, proteomic studies in nonmuscle cells have identified LAP3 in mitochondria-associated protein networks^20,21^. Furthermore, LAP3 may act as a mitochondrial protease involved in mitochondrial protein quality control in skeletal muscle, suggesting a role in the maintenance of mitochondrial homeostasis^22^. Because of its roles in mitochondrial activity and dynamics in skeletal muscle growth, fiber-type specification, metabolic adaptation, and endurance capacity^23^, LAP3 may contribute to skeletal muscle homeostasis through the regulation of mitochondrial function. The physiological role of LAP3 in skeletal muscle remains largely unclear in vivo. In particular, whether LAP3 regulates mitochondrial homeostasis and metabolic adaptation in skeletal muscle in vivo is unknown. To address this question, we generated LAP3-deficient mice and examined the role of LAP3 in skeletal muscle homeostasis with a particular focus on mitochondrial and metabolic regulation.

## Results

### Generation and validation of whole-body LAP3-deficient mice

We generated whole-body LAP3-deficient (LAP3-KO) mice by introducing a frameshift mutation through a cytosine insertion at position 181 in exon 2 of the Lap3 gene (Figure 1A). The loss of LAP3 protein expression across multiple tissues was confirmed by western blot analysis (Figure 1B).

**Figure 1.**
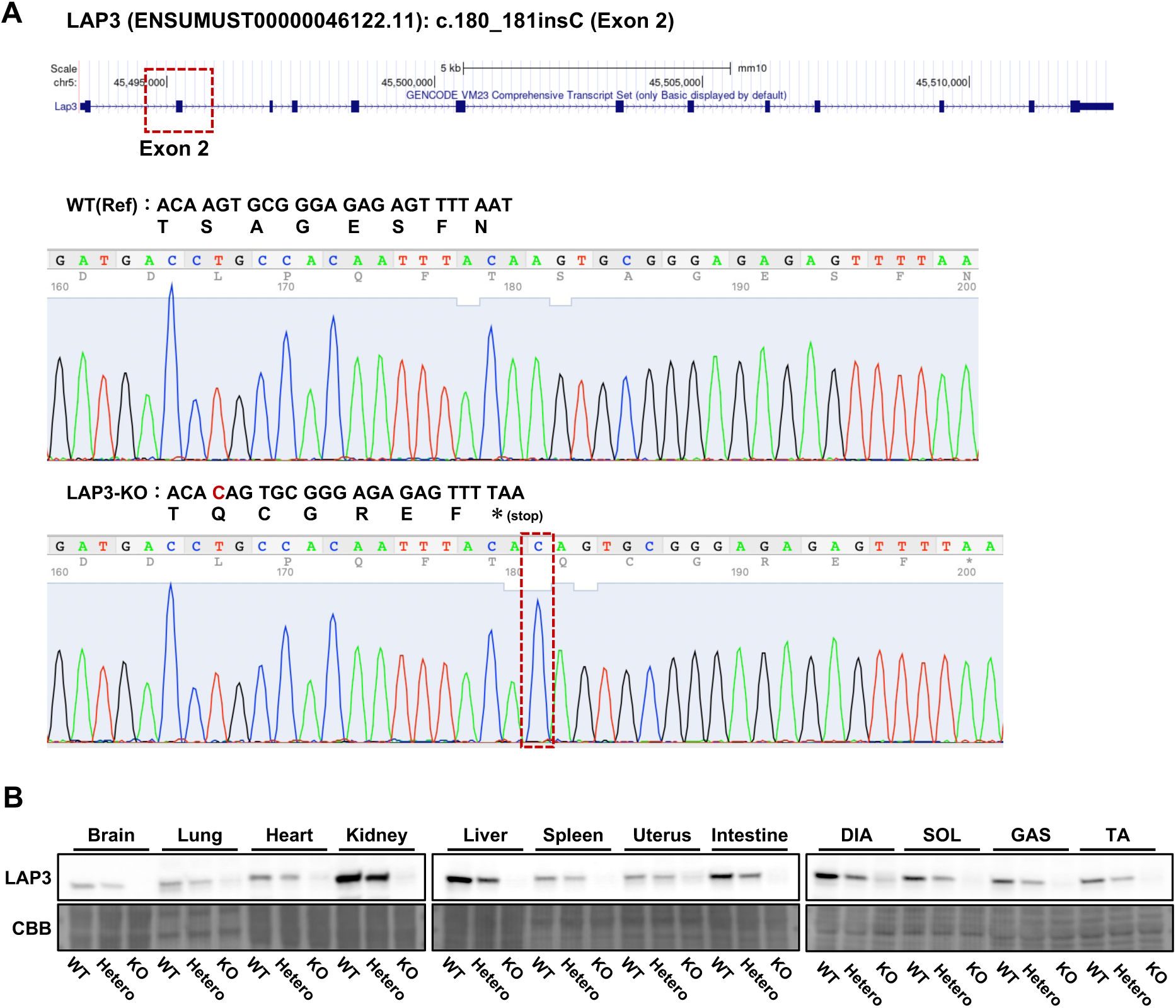
Generation and validation of whole-body LAP3-deficient mice. (A) Schematic representation of the constitutive *Lap3* knockout allele created by a cytosine insertion at nucleotide position 181 in exon 2, resulting in a frameshift mutation. (B) Representative western blots showing the absence of LAP3 protein expression in multiple tissues from LAP3-deficient (LAP3-KO) mice.

### LAP3 deficiency impairs skeletal muscle maintenance in a sex-dependent manner

Phenotypic characterization revealed that LAP3 deficiency reduced body weight in male and female mice (Figure 2A). Despite this reduction, no significant differences were observed in the absolute or body weight-normalized weights of major organs, including the heart, lung, liver, kidney, and spleen (Supplementary Figure 1), suggesting that the effects of LAP3 deficiency are not broadly distributed among tissues. In contrast, skeletal muscle was more prominently affected. The absolute weights of the tibialis anterior (TA) and gastrocnemius (GC) muscles were significantly decreased in female LAP3-KO mice (Figure 2B, C). These reductions were significant after normalization to body weight, indicating a disproportionate loss of TA and GC muscle mass relative to overall body weight. In contrast, soleus muscle weight was unchanged regardless of normalization (Figure 2D). In male mice, the absolute weights of the TA and GC muscles were also reduced by LAP3 deficiency; however, these differences were no longer evident following normalization to body weight (Figure 2E, F). Soleus muscle weight remained unchanged in males (Figure 2G). Taken together, these results indicate that LAP3 deficiency preferentially affects skeletal muscle maintenance rather than overall organ growth, and this phenotype is more pronounced in females. The preservation of soleus muscle weight further suggests that the effects of LAP3 deficiency are not uniform across all skeletal muscles.

**Figure 2.**
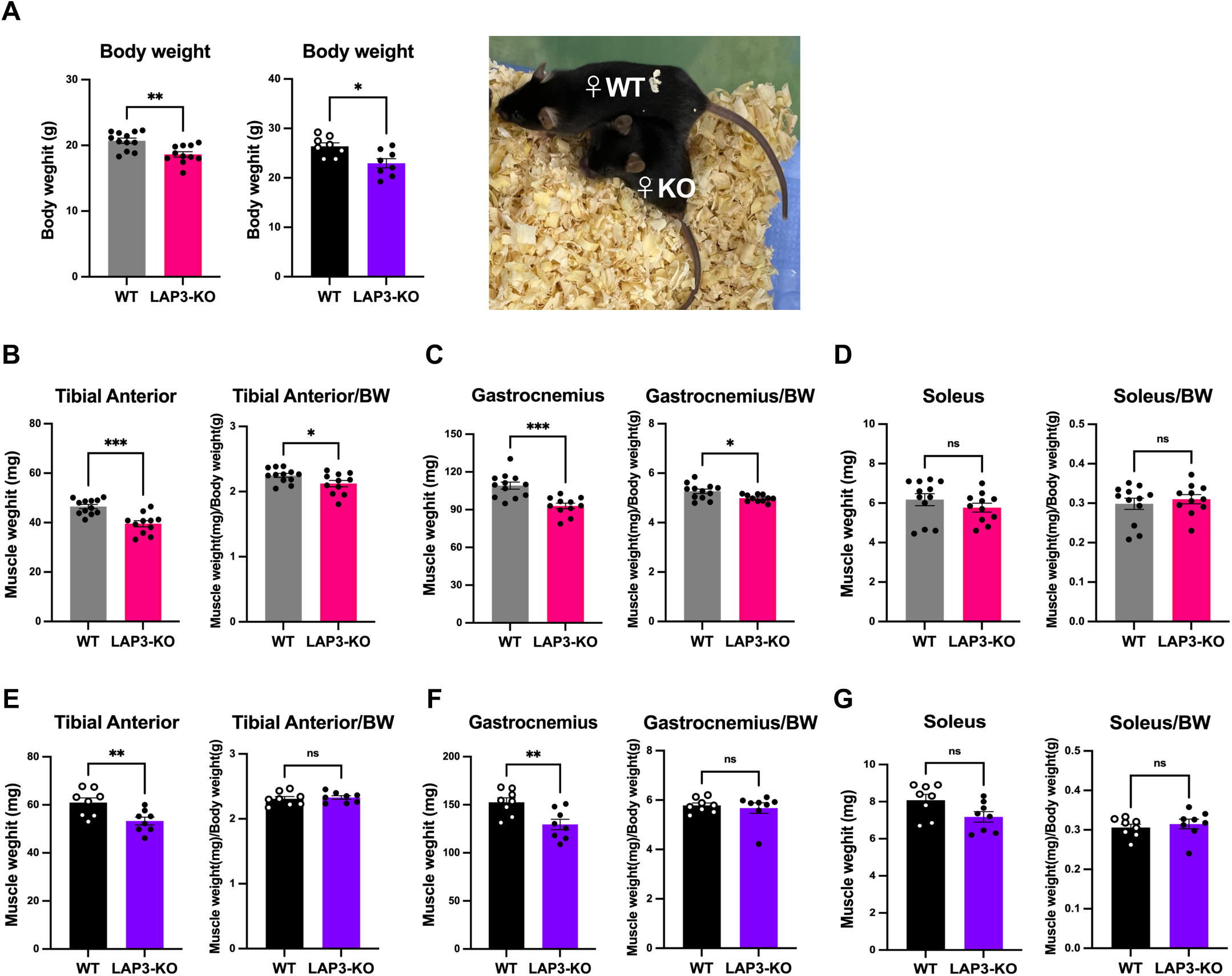
LAP3 deficiency disrupts skeletal muscle maintenance in a sex-dependent manner. (A) Body weight of female and male WT and LAP3-KO mice. (B–D) Absolute and body weight-normalized weights of the tibialis anterior (TA), gastrocnemius (GC), and soleus (SOL) muscles in female WT and LAP3-KO mice. (E–G) Corresponding analyses in male WT and LAP3-KO mice. The data are presented as the mean ± SEM (n = 8–12 mice per group). Statistical significance was assessed using an unpaired Student’s t-test. *P < 0.05, **P < 0.01, ***P < 0.001.

### LAP3 deficiency alters mitochondrial morphology, with female mice exhibiting greater metabolic vulnerability

Histological analysis of hematoxylin and eosin (H&E)-stained tibialis anterior (TA) sections showed no evidence of muscle damage, such as centralized nuclei or overt fiber degeneration, in either genotype (Figure 3A). Nevertheless, female LAP3-KO mice exhibited reduced myofiber size, as indicated by decreased Feret diameter and perimeter, together with a leftward shift in myofiber size distribution (Figure 3B, C). In contrast, no significant alterations in these parameters were observed in males (Figure 3D–F), suggesting that female skeletal muscle is more susceptible to LAP3 deficiency.

**Figure 3.**
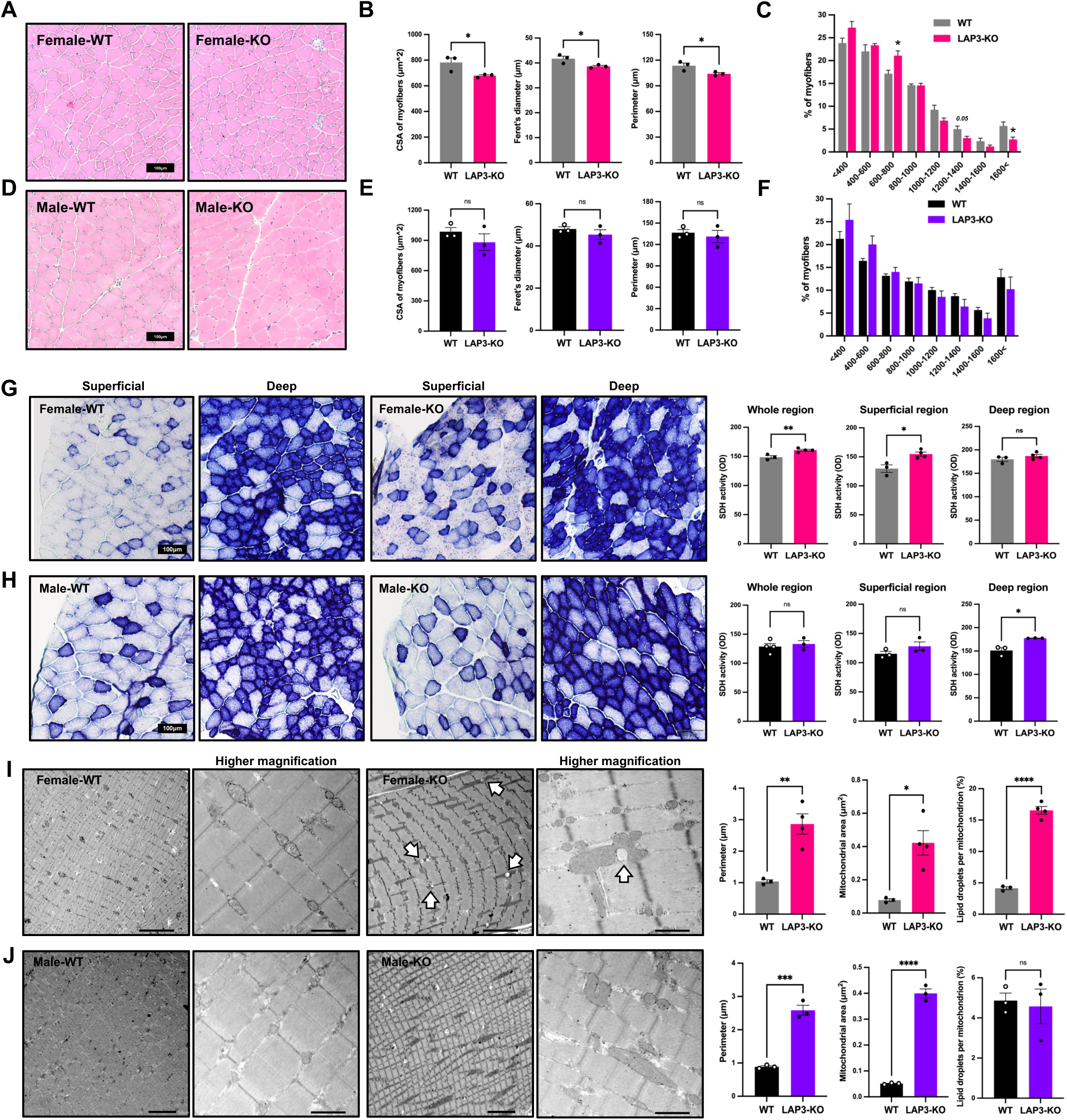
LAP3 deficiency alters mitochondrial morphology, with female mice exhibiting greater metabolic vulnerability. (A) Representative hematoxylin and eosin (H&E)-stained tibialis anterior (TA) muscle sections. (B, C) Quantification of myofiber size and distribution in female WT and LAP3-KO mice. (D–F) Corresponding analyses of TA myofiber size in male WT and LAP3-KO mice. (G) Succinate dehydrogenase (SDH) staining and quantification of oxidative enzyme activity in female TA muscle. (H) Corresponding SDH analyses in male TA muscle. (I, J) Representative transmission electron micrographs and quantification of mitochondrial perimeter and area in skeletal muscle. The white arrows indicate lipid droplets located adjacent to the mitochondria. The data are presented as the mean ± SEM (n = 3–4 mice per group). Statistical significance was assessed using an unpaired Student’s t-test. *P < 0.05, **P < 0.01, ***P < 0.001, ****P < 0.0001.

Succinate dehydrogenase (SDH) staining revealed sex-dependent changes in oxidative enzyme activity. Female LAP3-KO mice showed increased SDH activity throughout the muscle, with a particularly pronounced increase in the superficial region, whereas no significant changes were observed in the deep region (Figure 3G). In contrast, males showed no differences in the whole muscle or superficial region, but they exhibited increased SDH activity in the deep region (Figure 3H), which indicates sex-dependent alterations in oxidative metabolism.

Transmission electron microscopy revealed mitochondrial enlargement in skeletal muscle of both sexes. Quantitative analysis exhibited significant increases in mitochondrial perimeter and area in LAP3-KO mice compared with WT controls (Figure 3I, J). Notably, lipid droplet accumulation in proximity to mitochondria was observed exclusively in female LAP3-KO mice, whereas these changes were not evident in males, suggesting sex-dependent alterations in mitochondrial lipid handling. Taken together, these results indicate that LAP3 deficiency is associated with mitochondrial enlargement in skeletal muscle irrespective of sex, whereas female mice display additional alterations in oxidative metabolism and lipid handling.

### LAP3 deficiency is associated with altered mitochondrial characteristics and mitochondrial quality control in a sex-dependent manner

Functional analyses revealed that handgrip strength was preserved in both sexes, whereas endurance exercise capacity was significantly decreased only in female LAP3-KO mice (Figure 4A, B), which indicates a female-specific impairment in exercise performance. Although citrate synthase activity was unchanged in both sexes, normalization to mitochondrial density observed in electron microscopy images revealed reduced citrate synthase activity per mitochondrial content in female and male LAP3-KO mice (Figure 4C, D). Consistent with this observation, the expression of oxidative phosphorylation (OXPHOS) proteins, including Atp5a, Uqcrc2, Mtco1, Sdhb, and Ndufb8, was unchanged in both sexes (Figure 4E, F). Pgc1α expression also remained unchanged (Figure 4G, H), suggesting that mitochondrial enlargement was not attributable to enhanced mitochondrial biogenesis and may reflect altered mitochondrial homeostasis. Sex-dependent responses to LAP3 deficiency were observed in proteins involved in mitochondrial dynamics. In females, the expression of Mfn2, Opa1, and cytochrome c was increased, whereas Drp1 remained unchanged (Figure 4G), suggesting that enhanced mitochondrial fusion contributes to mitochondrial enlargement. In contrast, males tended towards increased cytochrome c expression, with no significant changes in Mfn2, Opa1, or Drp1 (Figure 4H). Taken together, these results indicate that LAP3 deficiency is associated with alterations in mitochondrial quality in both sexes, whereas female mice exhibit additional alterations in mitochondrial dynamics that are associated with reduced endurance capacity.

**Figure 4.**
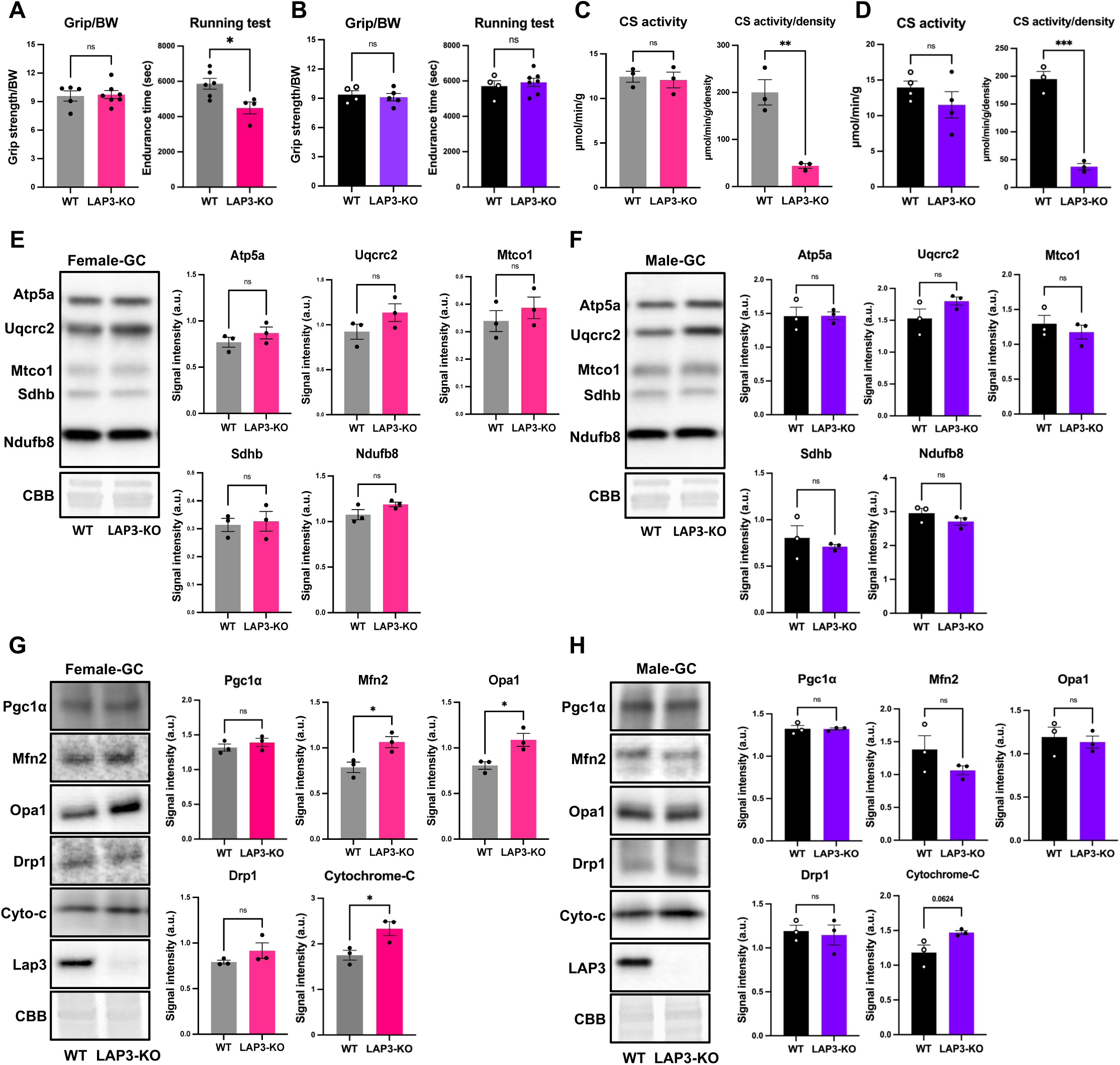
LAP3 deficiency is associated with altered mitochondrial characteristics and mitochondrial quality control in a sex-dependent manner. (A, B) Handgrip strength and endurance exercise capacity in female and male wild-type (WT) and LAP3-KO mice. (C, D) Citrate synthase activity and citrate synthase activity normalized to mitochondrial density estimated from electron microscopy images in female and male mice. (E, F) Representative western blots and quantification of the OXPHOS proteins, ATP5A, UQCRC2, MTCO1, SDHB, and NDUFB8, in female and male mice. (G, H) Representative western blots and quantification of PGC1α and proteins involved in mitochondrial dynamics, including MFN2, OPA1, DRP1, and cytochrome c, in female and male mice. The data are presented as the mean ± SEM (n = 3–7 mice per group). Statistical significance was assessed using an unpaired Student’s t-test. *P < 0.05, **P < 0.01, ***P < 0.001.

### LAP3 deficiency alters systemic energy metabolism and substrate utilization dynamics in female mice

To assess systemic metabolic function, we performed indirect calorimetry in LAP3-KO and wild-type mice. Female LAP3-KO mice exhibited significantly higher oxygen consumption (VOLJ), carbon dioxide production (VCOLJ), and energy expenditure compared with wild-type controls, which was accompanied by significant genotype and genotype-by-time interaction effects for each parameter (Figure 5A–C, left panels). Although no overall genotype effect was observed for the respiratory exchange ratio (RER), a significant genotype-by-time interaction was evident (Figure 5D, left panel). Similarly, lipid oxidation showed a significant genotype-by-time interaction together with a trend toward a genotype effect, whereas carbohydrate oxidation exhibited a significant genotype-by-time interaction without an overall genotype effect (Figure 5E, F, left panels). These results indicate altered diurnal dynamics of substrate utilization in female LAP3-KO mice.

**Figure 5.**
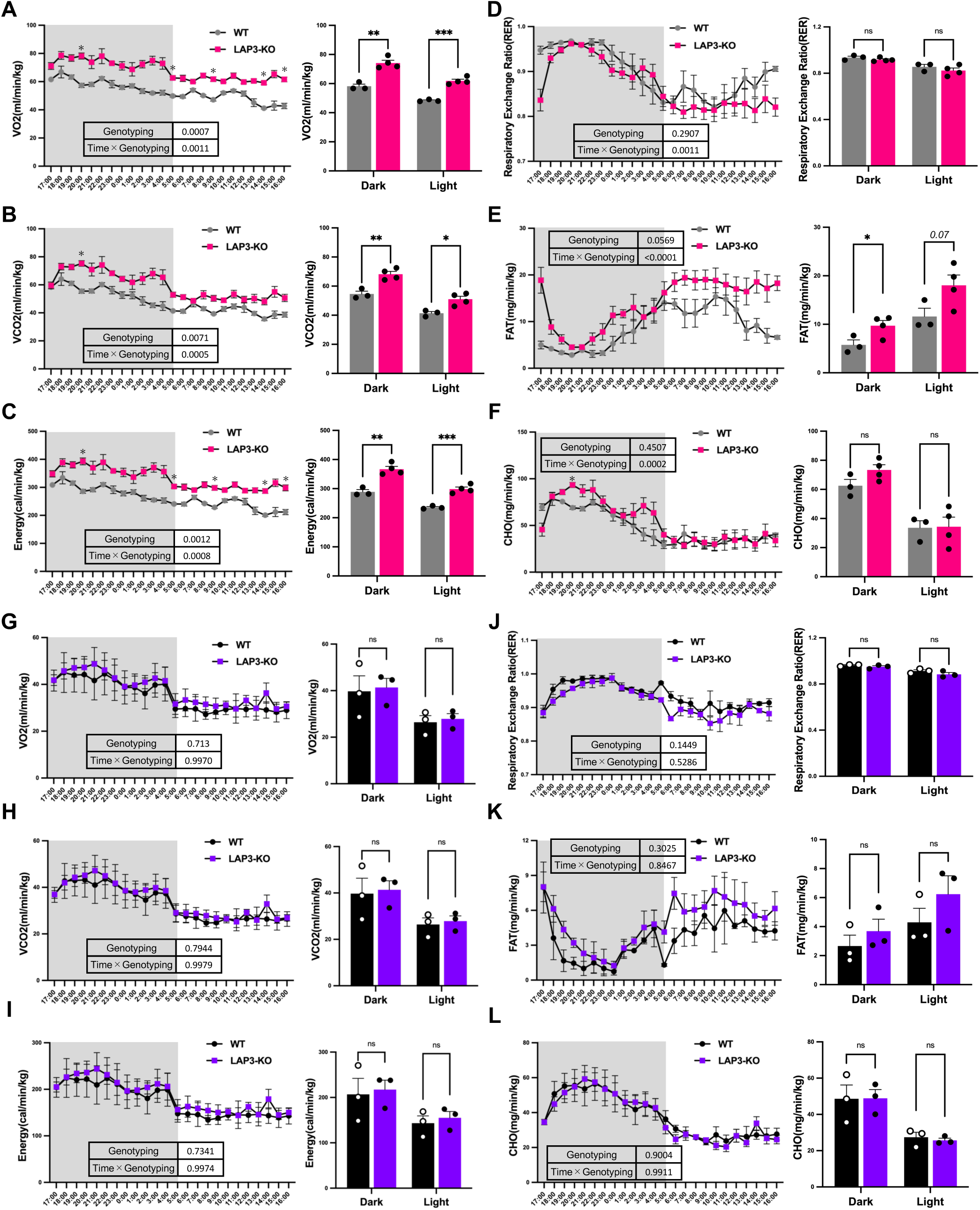
LAP3 deficiency alters systemic energy metabolism and substrate utilization dynamics in female mice. (A–F) Indirect calorimetry analysis of oxygen consumption (VOLJ), carbon dioxide production (VCOLJ), energy expenditure, respiratory exchange ratio (RER), lipid oxidation, and carbohydrate oxidation in female WT and LAP3-KO mice. (G–L) Corresponding analyses in male WT and LAP3-KO mice. In each panel, the left graph shows the 24-h profile of the indicated parameter, whereas the right graph shows the average values during the light and dark phases. The data are presented as the mean ± SEM (n = 3–4 mice per group). Statistical significance for the 24-h time-course data was assessed using a mixed-effects model (REML) with genotype and time (1-h bins) as factors, followed by Bonferroni’s multiple-comparisons test. Comparisons between genotypes within the light and dark phases were done using an unpaired Student’s t-test. *P < 0.05, **P < 0.01, ***P < 0.001.

To further characterize these metabolic alterations, the data were analyzed separately during the dark and light phases. Female LAP3-KO mice exhibited significantly increased VOLJ, VCOLJ, and energy expenditure during both phases (Figure 5A–C, right panels). Although RER and carbohydrate oxidation did not differ between genotypes, lipid oxidation was significantly increased during the dark phase and also tended to increase during the light phase in LAP3-KO females (Figure 5D–F, right panels).

In contrast, male LAP3-KO mice exhibited no significant alterations in metabolic parameters compared with the wild-type controls (Figure 5G–L). Food intake was unchanged between female wild-type and LAP3-KO mice (Supplementary Figure 2), indicating that the metabolic alterations were unlikely to be attributable to differences in energy intake. Taken together, these results indicate that LAP3 deficiency induces female-specific alterations in systemic energy metabolism and substrate utilization dynamics.

### LAP3 deficiency is associated with sex-dependent alterations in mitochondrial quality control and proteostatic pathways

Proteomic analyses revealed metabolic remodeling as a common feature of LAP3 deficiency. GO-BP analysis indicated enrichment of lipid metabolism-related pathways in both sexes, whereas males additionally exhibited enrichment of pathways associated with fiber-type transition (Figure 6A, B, Supplementary Figure 3A, B).

**Figure 6.**
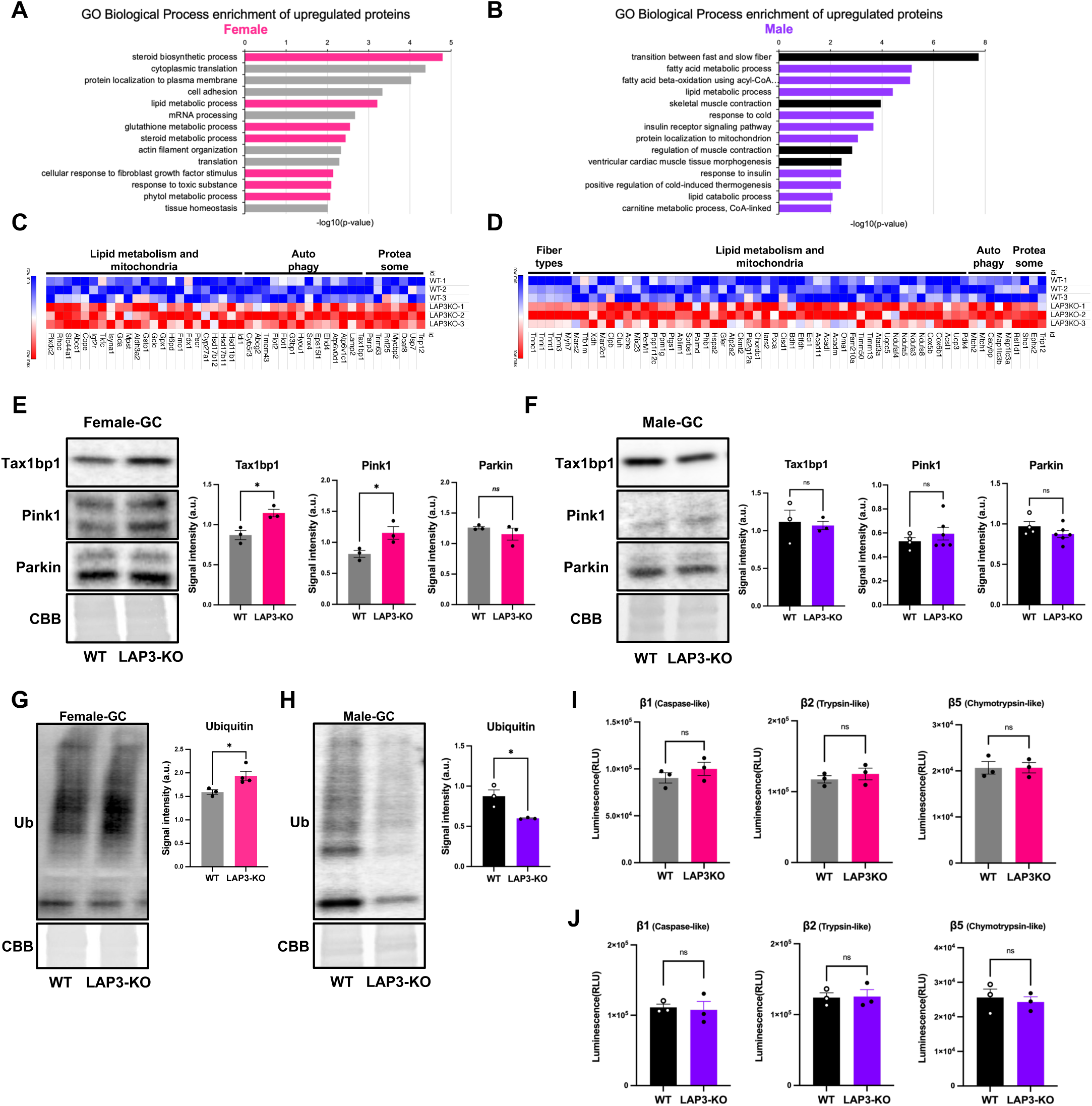
LAP3 deficiency alters pathways related to mitochondrial quality control and proteostasis in a sex-dependent manner. (A, B) Gene Ontology biological process (GO-BP) analysis showing the enriched pathways in female and male LAP3-KO skeletal muscle. (C, D) Representative proteins associated with lipid metabolism, mitochondrial function, proteostasis, and muscle fiber type were identified by proteomic analysis in females and males. (E, F) Representative western blots and quantification of TAX1BP1, PINK1, and PARKIN expression in female and male mice. (G, H) Representative western blots and quantification of ubiquitinated proteins in female and male mice. (I–K) Activities of the proteasomal catalytic subunits β1 (caspase-like), β2 (trypsin-like), and β5 (chymotrypsin-like) in female and male mice. The data are presented as the mean ± SEM (n = 3–4 mice per group). Statistical significance was assessed using an unpaired Student’s t-test. *P < 0.05, **P < 0.01, ***P < 0.001.

Because of the mitochondrial abnormalities observed in LAP3-KO mice, we evaluated markers related to mitochondrial quality control. In females, Tax1bp1 and Pink1 expression were significantly increased, whereas Parkin expression was unchanged (Figure 6E). In contrast, no significant changes were observed in males (Figure 6F), suggesting female-specific alterations in mitochondrial quality control pathways.

To further evaluate proteostatic regulation, we examined markers associated with autophagy and protein turnover. The expression of p62 and LC3 was unchanged in both sexes (Supplementary Figure 3C, D). In contrast, ubiquitinated proteins accumulated in female LAP3-KO mice, but were reduced in males (Figure 6G, H), indicating sex-dependent alterations in proteostatic regulation. Despite these changes, the activities of the proteasomal catalytic subunits β1, β2, and β5 were unchanged in either sex (Figure 6I, J), suggesting that LAP3 deficiency influences protein quality control independently of overt changes in proteasome catalytic activity.

Finally, male LAP3-KO mice showed increased expression of *Anpep* and *Tpp2* (Supplementary Figure 4), which suggests sex-dependent remodeling of aminopeptidase networks in response to LAP3 deficiency. Taken together, these results indicate that LAP3 deficiency induces metabolic remodeling and is associated with sex-specific alterations in mitochondrial quality control and proteostasis pathways in skeletal muscle.

### LAP3 modulates mitochondrial morphology in cultured myogenic cells

To examine the relationship between LAP3 and mitochondria in myogenic cells, we confirmed LAP3 expression in cytosolic and mitochondrial fractions of C2C12 myoblasts (Figure 7A). LAP3 knockdown in C2C12 myotubes increased Mitobright fluorescence intensity normalized to the myotube area (Figure 7B). In contrast, LAP3 overexpression in C2C12 myoblasts induced mitochondrial fragmentation, as assessed by Mitotracker staining (Figure 7C). These results support a role for LAP3 in the regulation of mitochondrial morphology.

**Figure 7.**
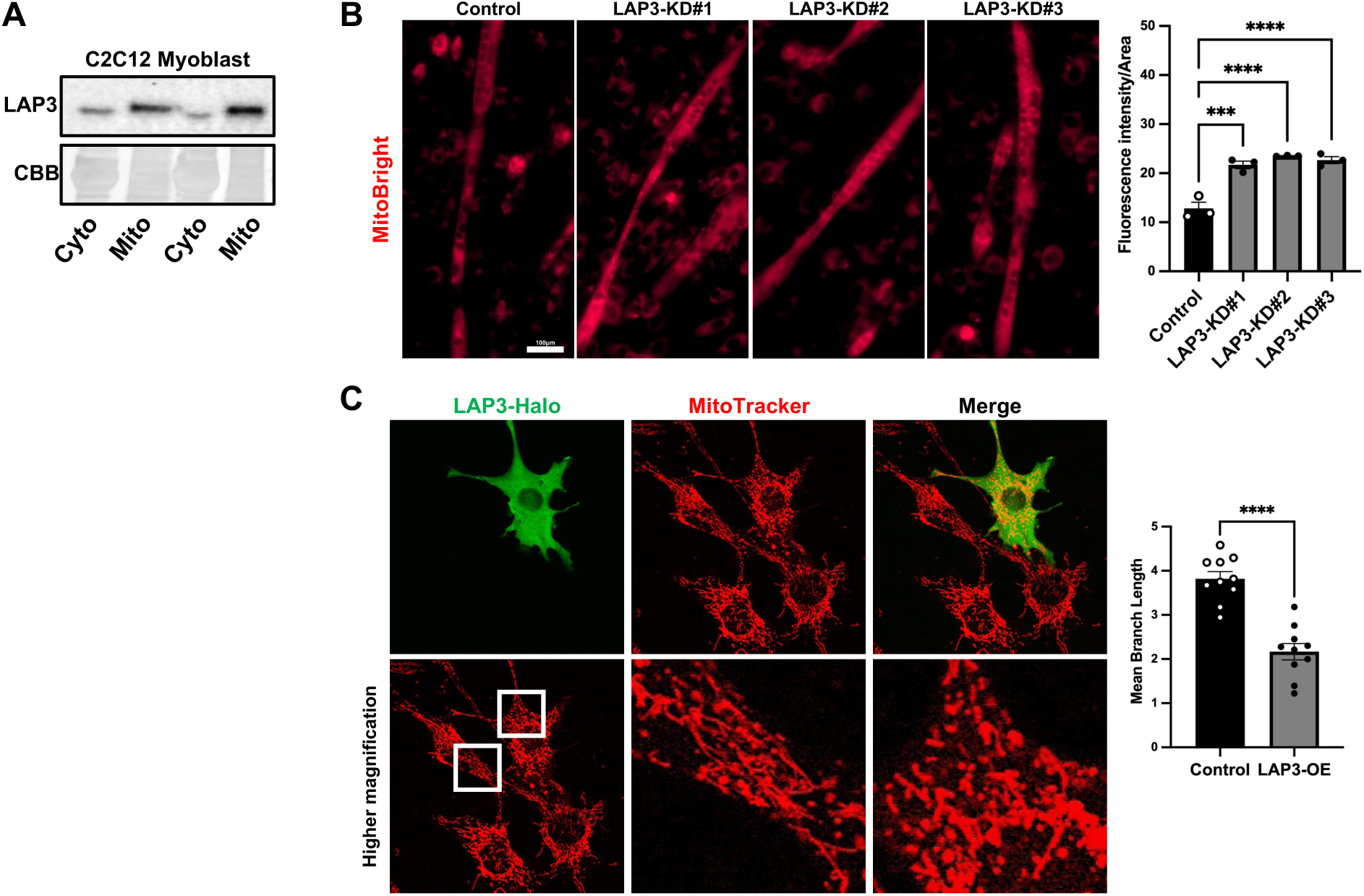
LAP3 modulates mitochondrial morphology in cultured myogenic cells. (A) LAP3 expression in cytosolic and mitochondrial fractions of C2C12 myoblasts. (B) Representative MitoBright staining and quantification of MitoBright fluorescence intensity normalized to myotube area in LAP3-knockdown (KD) C2C12 myotubes. Approximately 10 myotubes were analyzed per well, and the mean value from each well was used for statistical analysis (n = 3 wells per group). (C) Representative MitoTracker staining of LAP3-overexpressing C2C12 myoblasts showing altered mitochondrial morphology. The data are presented as the mean ± SEM. Statistical significance was assessed using a one-way ANOVA followed by Dunnett’s multiple-comparisons test for panel B and an unpaired Student’s t-test for panel C. *P < 0.05, **P < 0.01, ***P < 0.001.

## Discussion

LAP3 has been implicated in myogenic differentiation; however, its physiological role in mature skeletal muscle remains unclear. In the present study, we showed that loss of LAP3 disrupts skeletal muscle homeostasis, resulting in mitochondrial remodeling, altered metabolic adaptation, and impaired exercise capacity. These results identify LAP3 as a previously unrecognized regulator of skeletal muscle integrity and extend its functional significance beyond myogenic regulation.

Previous studies identified LAP3 within mitochondria-associated protein networks and hypothesized that LAP3 functions as a mitochondrial protease involved in mitochondrial protein quality control^20–22^. Consistent with this possibility, LAP3 was detected in mitochondrial fractions of C2C12 cells, and attenuating LAP3 expression alters mitochondrial morphology, which supports a role for LAP3 in the regulation of mitochondrial homeostasis. In the present study, LAP3 deficiency induced marked mitochondrial enlargement in skeletal muscle of both sexes. This was accompanied by reduced citrate synthase activity relative to mitochondrial content, whereas expression of OXPHOS proteins and Pgc1α remained unchanged. These results indicate that mitochondrial enlargement is unlikely to result from enhanced mitochondrial biogenesis. Instead, the enlarged mitochondria likely reflect altered mitochondrial homeostasis and remodeling. Similar mitochondrial enlargement has been observed in conditions of disrupted fusion–fission balance or impaired mitochondrial turnover^24,25^. In female LAP3-KO mice, increased Mfn2 and Opa1 expression further suggests that enhanced mitochondrial fusion may contribute to this phenotype. Notably, the reduction in citrate synthase activity relative to mitochondrial content suggests that mitochondrial enlargement is not accompanied by a proportional increase in oxidative capacity. Thus, the enlarged mitochondria observed in LAP3-deficient skeletal muscle may represent a qualitatively altered mitochondrial population rather than a simple expansion of mitochondrial mass. Taken together, these findings support a model in which LAP3 contributes to the maintenance of mitochondrial homeostasis in skeletal muscle, whereas its loss promotes mitochondrial remodeling associated with altered mitochondrial quality and function.

Female LAP3-KO mice exhibited increased oxygen consumption, carbon dioxide production, and energy expenditure, together with altered substrate utilization dynamics. Notably, lipid oxidation was increased during the dark phase, whereas lipid droplets accumulated around the mitochondria in skeletal muscle. These observations suggest that LAP3 deficiency alters the metabolic response to energetic demand. Skeletal muscle in females is generally characterized by greater mitochondrial abundance and a stronger reliance on lipid metabolism compared with that of males^26–31^. Therefore, disturbances in mitochondrial homeostasis may have a greater physiological impact on females. Interestingly, mitochondrial enlargement was observed in both sexes, whereas the metabolic and functional consequences were largely restricted to females. This dissociation suggests that mitochondrial remodeling itself may not be sufficient to cause overt dysfunction, and the capacity to adapt to such remodeling may be a key determinant of phenotypic severity. The coexistence of increased energy expenditure altered substrate utilization, lipid droplet accumulation, and reduced endurance capacity suggests that LAP3 deficiency disrupts metabolic adaptation in female skeletal muscle. Moreover, the accumulation of lipid droplets around mitochondria despite increased lipid oxidation suggests a mismatch between lipid availability and utilization in LAP3-deficient skeletal muscle. Such a mismatch may contribute to metabolic stress and reduced exercise performance^32–34^. Taken together, these results support the concept that LAP3 contributes to the maintenance of metabolic flexibility by preserving mitochondrial homeostasis, and its disruption preferentially affects female skeletal muscle.

Because LAP3 functions downstream of the ubiquitin–proteasome system as a post-proteolytic aminopeptidase, its deficiency may affect cellular homeostasis through mechanisms distinct from canonical protein degradation pathways^18^. In the present study, female LAP3-KO mice showed increased expression of Pink1 and Tax1bp, together with the accumulation of ubiquitinated proteins, whereas proteasome activity, p62 expression, and LC3 expression remained unchanged. These results suggest that LAP3 deficiency is associated with selective alterations in mitochondrial quality control and proteostatic regulation rather than global activation of autophagy or impairment of proteasome catalytic function. The persistence of mitochondrial enlargement despite increased expression of mitochondrial quality control-associated markers suggests that these responses may represent compensatory adaptations to mitochondrial stress, rather than effective mitochondrial clearance^35^. Because LAP3 acts at the final stage of proteasome-dependent peptide degradation, its disruption may alter intracellular peptide homeostasis, thereby linking post-proteolytic peptide processing to mitochondrial homeostasis^36^. In addition, previous studies have indicated that LAP3 functions as a mitochondrial protease involved in mitochondrial protein quality control^20–22^. Therefore, the mitochondrial abnormalities observed in LAP3-deficient skeletal muscle may not only reflect altered peptide homeostasis, but also impaired mitochondrial proteostasis. Further studies are needed to determine whether LAP3 directly regulates mitochondrial protein turnover and quality control pathways in skeletal muscle.

Male LAP3-KO mice exhibited increased expression of the aminopeptidases, *Anpep* and *Tpp2,* despite relatively mild physiological phenotypes. Because of the role of aminopeptidases in peptide turnover and amino acid homeostasis, these changes may represent compensatory adaptations that partially offset the loss of LAP3 function. Notably, TPP2 has been implicated in the maintenance of metabolic, lysosomal, and cellular homeostasis beyond its canonical role in peptide degradation^37,38^, which suggests that increased *Tpp2* expression reflects a broader adaptive response to proteostatic stress. Although further studies are needed to establish the functional significance of these changes, the present findings suggest that males and females engage distinct adaptive responses to the disruption of LAP3-dependent homeostatic pathways.

This study had several limitations that should be acknowledged. First, because a constitutive whole-body LAP3 knockout model was used, the observed phenotypes may reflect both muscle-intrinsic functions of LAP3 and systemic or developmental effects arising from lifelong LAP3 deficiency. Second, although our findings support a link between LAP3 and mitochondrial homeostasis, mitochondrial respiratory function was not directly assessed and requires future study. Finally, the mechanisms underlying the pronounced sexual dimorphism observed in LAP3-KO mice remain unclear, as the contribution of sex hormone signaling was not directly examined. Addressing these limitations will clarify how LAP3 regulates skeletal muscle homeostasis. Figure 8 summarizes the proposed role of LAP3 in maintaining skeletal muscle mitochondrial homeostasis and the sex-dependent consequences of LAP3 deficiency.

**Figure 8.**
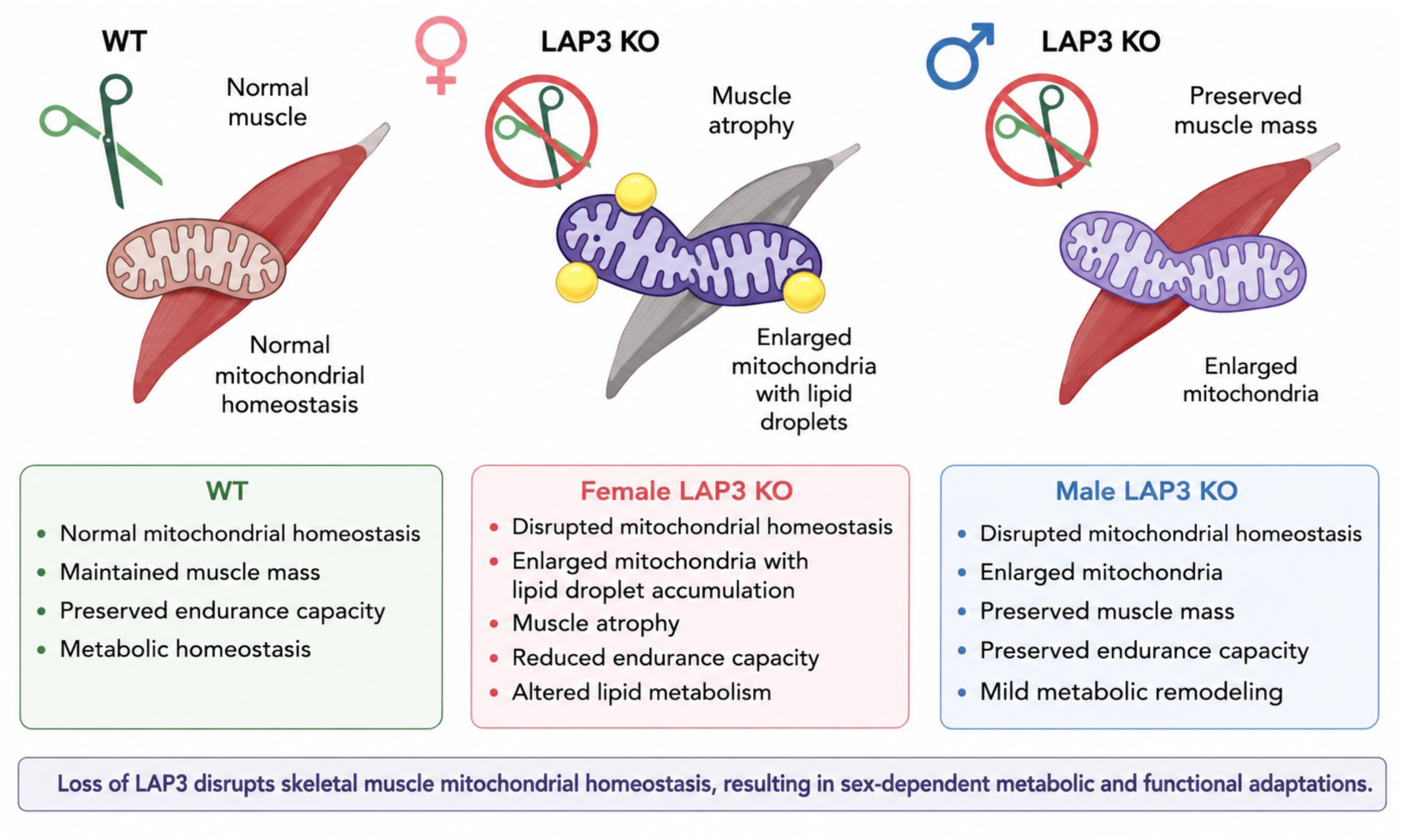
Proposed model of LAP3-dependent regulation of skeletal muscle mitochondrial homeostasis. Under physiological conditions, LAP3 plays a crucial role in maintaining mitochondrial homeostasis, metabolic flexibility, and skeletal muscle integrity. Loss of LAP3 induces mitochondrial enlargement in both female and male skeletal muscle, indicating disrupted mitochondrial homeostasis. In female LAP3-KO mice, mitochondrial remodeling is associated with lipid droplet accumulation, muscle atrophy, reduced endurance capacity, and altered lipid metabolism. In contrast, male LAP3-KO mice exhibit mitochondrial enlargement with preserved muscle mass and endurance capacity, accompanied by relatively mild metabolic remodeling. These findings support a role for LAP3 in maintaining skeletal muscle mitochondrial homeostasis and highlight sex-dependent physiological responses to LAP3 deficiency.

In conclusion, we identified LAP3 as a previously unrecognized regulator of skeletal muscle homeostasis. LAP3 deficiency is associated with altered mitochondrial morphology, metabolic adaptation, and proteostatic responses, resulting in a predominantly female muscle phenotype. Taken together, these results provide insight into the mechanisms that maintain skeletal muscle integrity and support a role for LAP3 in the regulation of mitochondrial homeostasis.

## Methods

### Generation of LAP3 knockout mice

Lap3 knockout mice were generated by CRISPR/Cas9-mediated genome editing (SetsuroTech, Tokushima, Japan). Guide RNAs were designed to target exon 2 of the Lap3 gene, resulting in the introduction of a single cytosine insertion at nucleotide position 181 and the generation of a frameshift mutation. Edited zygotes were transferred into pseudopregnant recipient females to generate founder (F0) mice. Founder mice were crossed with C57BL/6J mice (Charles River Laboratories, Kanagawa, Japan) to obtain heterozygous offspring, which were subsequently used to establish the colony. Homozygous Lap3-deficient (Lap3-KO) mice were generated by breeding heterozygous or homozygous mice, and wild-type littermates were used as controls. Genomic DNA was isolated from tail biopsies, and the genotype were confirmed by PCR followed by Sanger sequencing. Amplification was performed using the primers Lap3LJF (5′LJCCTCTGCGGAAAAAGACAGTLJ3′) and Lap3LJR (5′LJCCAGAACTCCAGGGAATGAGLJ3′), yielding a 400-bp PCR product.

### Experimental animals

Wild-type (WT) and Lap3-knockout (LAP3-KO) mice on a C57BL/6J background were used in this study. Male and female mice aged 3–4 months were analyzed separately. All mice were housed in a temperature-controlled facility (22–24 °C) under a 12-h light/12-h dark cycle with ad libitum access to standard chow and water. All experimental procedures were approved by the Institutional Animal Care and Use Committee of the University of Electro-Communications (Protocols No. A40 and G2-1003) and were conducted in accordance with institutional guidelines and national regulations for the care and use of laboratory animals.

### Cell culture and differentiation

Mouse C2C12 myoblast cell lines (Cell No. RCB0987, RIKEN BioResource Research Center, Tsukuba, Japan) were maintained in Dulbecco’s modified Eagle’s medium (DMEM; FUJIFILM Wako Pure Chemical Corporation, Osaka, Japan) supplemented with 10% fetal bovine serum (Thermo Fisher Scientific, Waltham, MA, USA) and 1% penicillin–streptomycin (FUJIFILM Wako) at 37 °C in a humidified atmosphere of 5% COLJ. For myogenic differentiation, confluent C2C12 myoblasts were transfected with small interfering RNA targeting Lap3 (siLap3) or non-targeting control siRNA (Sigma-Aldrich, Merck KGaA, Darmstadt, Germany) using Lipofectamine RNAiMAX (Thermo Fisher Scientific) according to the manufacturer’s instructions. Twenty-four hours after transfection, the culture medium was replaced with differentiation medium (DMEM supplemented with 2% horse serum [Thermo Fisher Scientific] and 1% penicillin–streptomycin [FUJIFILM Wako]), and cells were induced to differentiate into myotubes for 7 days. The differentiation medium was changed every 2 days throughout the differentiation period.

For LAP3 overexpression experiments, C2C12 myoblasts were transiently transfected with a C-terminal HaloTag-fused LAP3 expression plasmid (pFC14K-LAP3) generated using the pFC14K HaloTag CMV Flexi Vector (Promega, Madison, WI, USA) using Lipofectamine 3000 (Thermo Fisher Scientific). After incubation with the transfection mixture for 6 h, the medium was replaced with fresh growth medium, and cells were cultured for an additional 18 h. Mitochondrial imaging and morphological analyses were subsequently performed in siRNA-treated myotubes and LAP3-overexpressing myoblasts.

### Mitochondrial staining and fluorescence analysis

On day 7 of myogenic differentiation, C2C12 myotubes were stained with MitoBright MT11 (Dojindo Technologies, Kumamoto, Japan) to visualize mitochondria. Cells were incubated with 0.1 μM MitoBright for 30 min at 37 °C, washed with PBS (FUJIFILM Wako), fixed with 4% paraformaldehyde (FUJIFILM Wako), and mounted with VECTASHIELD containing DAPI (Vector Laboratories, Newark, CA, USA). Fluorescence images were acquired using an Olympus fluorescence microscope (Evident Corporation, Tokyo, Japan). MitoBright fluorescence intensity was quantified using Fiji/ImageJ software (NIH) and normalized to myotube area. Approximately 10 myotubes were analyzed per well, and the mean value from each well was used for statistical analysis (n = 3 wells per group). For LAP3 overexpression experiments, cells were stained with Oregon Green HaloTag Ligand (Promega, Madison, WI, USA) and MitoTracker Red CMXRos (Thermo Fisher Scientific) for 20 min at 37 °C and imaged using an Olympus FV1000 confocal laser-scanning microscope (Olympus, Tokyo, Japan). Mitochondrial morphology was analyzed using the Mitochondria Analyzer plugin in Fiji/ImageJ, and mean branch length was used as an index of mitochondrial network morphology. Mitochondrial morphology was quantified in approximately 10 cells per group collected from several independent experiments. Mean branch length was calculated for each cell and used for statistical analysis.

### Exercise performance test

Grip strength was measured using a grip strength meter (Melquest, Toyama, Japan). Mice were allowed to grasp a wire mesh with their forelimbs, and their tails were gently pulled backward until they released their grip. Three measurements were obtained for each mouse, and the average value was used for analysis. Grip strength was normalized to body weight.

Endurance exercise capacity was assessed using a motorized treadmill system for small animals (Melquest). Mice were first acclimated by running at 10 m/min for 5 min. The endurance test then began at 10 m/min for 30 min, after which the speed was increased by 2 m/min every 15 min until exhaustion. Exhaustion was defined as the inability to resume running after remaining on the shock grid despite repeated mechanical or electrical stimulation^39^. Total running time was recorded as an index of endurance capacity.

### Mitochondrial enzyme activity

The maximal activities of citrate synthase (CS) were determined in muscle homogenates. In brief, the gastrocnemius was homogenized in 100 (vol/wt) of 100-mmol/L potassium phosphate buffer. Maximal activities were quantified using a microplate reader, as described previously^40^. CS activity was normalized to mitochondrial density quantified from electron microscopy images to estimate enzymatic activity relative to mitochondrial abundance.

### Indirect calorimetry and energy metabolism

Energy expenditure and substrate utilization were assessed using an indirect calorimetry system (Arco system, Chiba, Japan). Mice were housed individually in metabolic chambers (W120 × H150 × D240 mm) with free access to standard chow and water. Mice were acclimated to the chambers for 5–7 days until behavioral and metabolic parameters stabilized. After acclimation, oxygen consumption (VOLJ), carbon dioxide production (VCOLJ), and respiratory exchange ratio (RER) were continuously monitored for 24 h and recorded at 10-min intervals. Energy expenditure, carbohydrate (CHO) oxidation, and fat (FAT) oxidation were calculated from VOLJ and VCOLJ according to standard indirect calorimetry equations. Data were analyzed as (1) 24-h time-course plots using 1-h bin averages and (2) average values during the dark phase (17:00–05:00) and light phase (05:00–17:00).

### Tissue collection and processing

Immediately after sacrifice, body weight and the wet weights of the heart, lungs, liver, spleen, kidneys, tibialis anterior (TA), gastrocnemius (GC), and soleus muscles were measured. The TA and GC muscles were rapidly frozen in liquid nitrogen for subsequent biochemical analyses and stored at −80LJ°C. For histological analyses, TA and GC muscles were embedded in optimal cutting temperature (OCT) compound (Sakura Finetek USA, Torrance, CA, USA), frozen in isopentane precooled with liquid nitrogen, and then stored at −80LJ°C until cryosectioning.

### Hematoxylin and eosin (H&E) staining

For histological evaluation, cryosections (10 μm thickness) of tibialis anterior muscles were prepared using a Leica CM1950 cryostat (Leica, Nussloch, Germany). Sections were stained with hematoxylin (FUJIFILM Wako) for 10 min, followed by incubation in an alkaline solution (2% w/v magnesium sulfate and 0.2% w/v potassium bicarbonate in distilled water) for 15 min to enhance nuclear contrast. The sections were then counterstained with eosin (FUJIFILM Wako) for 2 min. After staining, the slides were dehydrated through a graded ethanol series (70% ethanol for 2 min, 80% ethanol for 2 min, 90% ethanol for 2 min, and 100% ethanol for 2 min), cleared in xylene for 2 min, and mounted under a coverslip with MountLJQuick mounting medium (Daido Sangyo, Tokyo, Japan). Stained sections were observed under a Nikon microscope for assessment of general muscle morphology. Tibialis anterior muscle size and individual myofiber size were quantified according to a previously described method using ImageJ-based image analysis^41^. Myofiber size was evaluated by measuring Feret diameter and perimeter, and myofiber size distribution profiles were generated from individual myofiber measurements.

### Succinate dehydrogenase (SDH) staining

For histochemical evaluation of oxidative capacity, cryosections (10 μm thickness) of tibialis anterior muscles were prepared. Sections were incubated in an SDH staining solution (FUJIFILM Wako) containing 0.2 M phosphate buffer (pH 7.5), 0.2 M sodium succinate, and 1 mg/mL nitroblue tetrazolium at 37LJ°C for 45 min. The staining solution was prepared fresh on the day of the experiment. After incubation, the slides were rinsed with distilled water, dehydrated through a graded ethanol series (70% ethanol for 2 min, 80% ethanol for 2 min, 90% ethanol for 2 min, and 100% ethanol for 2 min), cleared in xylene for 2 min, and mounted under a coverslip with MountLJQuick mounting medium (Daido Sangyo). SDHLJstained sections were observed under a Nikon microscope for assessment of oxidative fiber distribution.

### Transmission electron microscopy

The collected samples were preLJfixed with 2.5% glutaraldehyde in 0.1 M cacodylate buffer (pH 7.4) and postLJfixed with 1% osmium tetroxide. After dehydration in a graded ethanol series, the samples were embedded in epoxy resin. Transmission electron microscopy (TEM) was conducted to examine the morphological features of mitochondria in muscle in detail. TEM images were acquired using a JEMLJ1400 Flash transmission electron microscope (JEOL, Tokyo, Japan) operated at an accelerating voltage of 80 kV. Using the obtained images, the mitochondrial perimeter and area (crossLJsectional area) were measured for individual mitochondria. All tracings were performed by blinded researchers to avoid bias. Morphological indices were calculated using Fiji image analysis software (NIH).

### Western blotting analysis

Muscle tissues were homogenized in RIPA buffer supplemented with protease and phosphatase inhibitors (ATTO Corporation, Tokyo, Japan). Protein concentrations were determined using a BCA protein assay kit (Thermo Fisher Scientific), and samples were adjusted to equal protein amounts. After addition of sample buffer, samples for OXPHOS analysis were incubated at 50 °C for 15 min, whereas samples for all other targets were heated at 95 °C for 5 min. Proteins were then separated by SDS–PAGE using 4–20% gradient MiniLJPROTEAN TGX precast protein gels (BioLJRad, Hercules, CA, USA) and transferred to PVDF membranes using the Trans-Blot Turbo Transfer System (Bio-Rad). TMembranes were blocked with Bullet Blocking One (Nacalai Tesque, Kyoto, Japan) according to the manufacturer’s instructions. For OXPHOS proteins, membranes were incubated with primary antibodies for 1 h at room temperature with gentle agitation. For all other target proteins, membranes were incubated with primary antibodies overnight at 4 °C. Membranes were washed and incubated with horseradish peroxidase-conjugated secondary antibodies for 1 h at room temperature. Immunoreactive bands were detected using enhanced chemiluminescence reagents (Cytiva, Tokyo, Japan) and visualized with the VuLJC Chemiluminescence Imaging System (PopLJBio Imaging, Cambridge, UK). Total protein staining with Coomassie Brilliant Blue (ATTO Corporation, Tokyo, Japan) was used as a loading control. Band intensities were quantified using ImageJ software (NIH, Bethesda, MD, USA). The antibodies used in this study are listed in Supplementary Table 1.

### Proteomic sample preparation and nanoLC-MS/MS analysis

Muscle tissues were homogenized in 70% ethanol–0.5% acetic acid with zirconia beads, and the homogenate was centrifuged at 15,000 × g at 4 °C for 15 min. The pellet was solubilized in 100 mM Tris–HCl (pH 8.0), 4% SDS, and 20 mM NaCl using ultrasonication, and the protein concentration was adjusted to 75 ng/μL. Proteins were processed by SP3LJbased purification and digestion using SeraLJMag SpeedBeads (Cytiva) and Trypsin/LysLJC Mix (Promega, Madison, WI, USA), followed by reduction/alkylation, desalting, and resuspension. NanoLCLJMS/MS analysis was performed as a contract service by Kazusa DNA Research Institute (Kisarazu, Japan) using their “Kazusa DIA Standard” service. Peptides were loaded onto a 75 μm i.d. × 300 mm C18 column (1.7 μm, 100 Å, CoAnn Technologies) at 50 °C, with gradient separation using 0.1% formic acid in water and 80% acetonitrile. The data were processed using DIALJNN, and proteins passing 1% precursor-level and 1% protein-level FDR thresholds were used for downstream analysis. The identified proteins were further analyzed using DAVID for Gene Ontology (GO) and Kyoto Encyclopedia of Genes and Genomes (KEGG) pathway enrichment analyses.

### RNA isolation and quantitative real-time PCR

Total RNA was isolated from skeletal muscle tissues using the RNeasy Fibrous Tissue Mini Kit (QIAGEN, Hilden, Germany) according to the manufacturer’s instructions. cDNA was synthesized from the purified RNA using the QuantiTect Reverse Transcription Kit (QIAGEN), and the resulting cDNA was used as template for quantitative realLJtime PCR. qPCR reactions were performed using the SYBR Green qPCR Master Mix (Thermo Fisher Scientific) on an Applied Biosystems StepOne Real-Time PCR System (Thermo Fisher Scientific). All qPCR runs were performed using the instrument’s default settings, and relative gene expression levels were determined using the ΔΔCt method normalized to an appropriate housekeeping gene. All primer sequences are listed in Supplementary Table 2.

### Proteasome activity assay

Proteasomal activity was measured from skeletal muscle lysates using the Proteasome-Glo Assay (Promega) according to the manufacturer’s instructions. Skeletal muscle tissues were homogenized, and protein extracts were prepared using Pierce RIPA Buffer (Thermo Fisher Scientific). Protein concentration was adjusted to 0.1 μg/μL, consistent with the conditions used for Western blotting. Equal volumes of protein extract and Proteasome-Glo Reagent were mixed in a white 96LJwell plate, and the plates were incubated at room temperature in the dark for 30 min. Luminescence was measured using a GloMax Navigator System (Promega), and proteasome activity was expressed as relative luminescence units (RLU) normalized to total protein amount.

### Mitochondrial and cytoplasmic fractionation

Mitochondrial and cytoplasmic fractions were isolated from C2C12 myoblasts using the Mitochondrial Isolation Kit (TCI Chemicals, Tokyo, Japan) according to the manufacturer’s instructions. Briefly, C2C12 cells were harvested, homogenized, and subjected to a series of differential centrifugation steps to obtain both mitochondrial and cytoplasmic fractions. Each fraction was then used for downstream analyses.

### Statistical analysis

Statistical analyses were performed using GraphPad Prism 9 (GraphPad Software, San Diego, CA, USA). Comparisons between WT and LAP3-KO mice were performed using unpaired two-tailed Student’s t-tests. For indirect calorimetry time-course data collected at 24 time points (1-h intervals over 24 h), a mixed-effects model (REML) was used with genotype and time as fixed effects, followed by Bonferroni’s multiple-comparisons test. For mitochondrial fluorescence intensity measurements in C2C12 myotubes, comparisons among four groups were performed using one-way ANOVA followed by Dunnett’s multiple-comparisons test. Data are presented as mean ± SEM. All experiments were independently repeated two to three times to confirm reproducibility. Sample sizes (n) for each experiment are indicated in the corresponding figure legends. A value of P < 0.05 was considered statistically significant.

## Data availability

All data are available from the corresponding authors upon request. The proteomics datasets generated in this study have been deposited in the jPOST repository under accession number JPST004687. During the peer-review process, the datasets are available at https://repository.jpostdb.org/preview/3719625216a361b2a34017 using the access key: 8419. Processed proteomics data, including protein abundance values and differential expression analyses, are provided in the Supplementary table files. Source data are provided with this paper. All other data supporting the findings of this study are available from the corresponding authors upon reasonable request.

## Acknowledgements

We thank the members of our laboratories for their helpful discussions and technical support.

## Author contributions

S.O. conceived and designed the study, performed the experiments, analyzed and interpreted the data, prepared the figures, and wrote and revised the manuscript. R.M. and R.Na. contributed to animal experiments and data acquisition. A.T., R.K., K.B., and H.T. contributed to histological analyses and data acquisition. H.W. contributed to cell culture experiments. N.S., K.M., M.K., Y.Ki., M.S., and D.H. contributed to data interpretation and critically revised the manuscript. R.Ng. and Y.Ka. supervised the study, acquired funding, and edited and revised the manuscript. All authors reviewed and approved the final manuscript.

