## Supplementary Figures for "Leucine Aminopeptidase 3 Regulates Skeletal Muscle Mitochondrial Homeostasis with Sex-Dependent Metabolic Consequences"

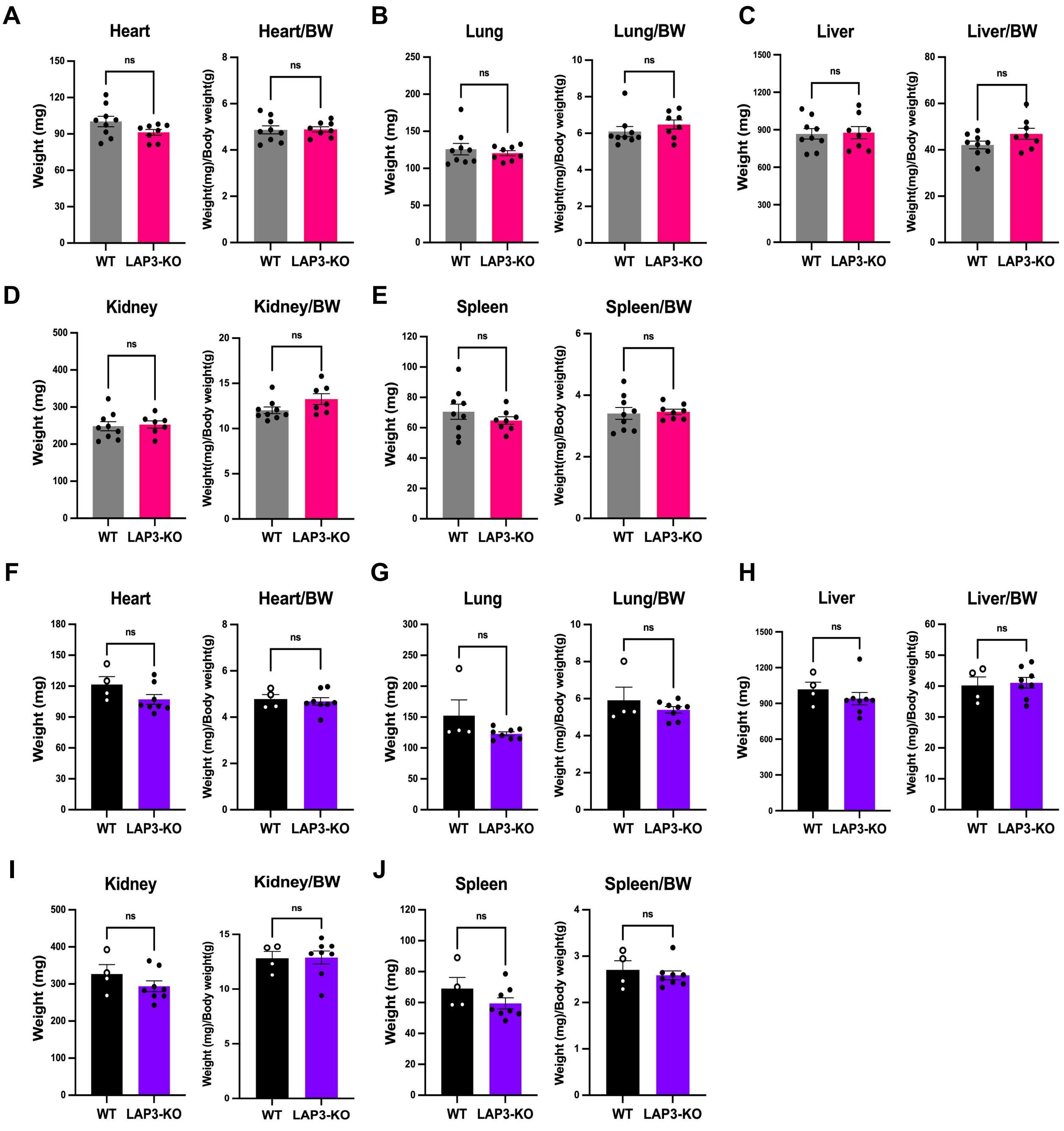


**Supplementary Figure 1. LAP3 deficiency does not markedly affect the weights of major organs**.

(A–E) Absolute and body-weight-normalized weights of the heart, lung, liver, kidney, and spleen in female WT (n = 9) and LAP3-KO (n = 8) mice. (F–J) Corresponding analyses in male WT (n = 4) and LAP3-KO (n = 8) mice. Data are presented as mean ± SEM. Statistical significance was assessed using an unpaired Student's t-test.

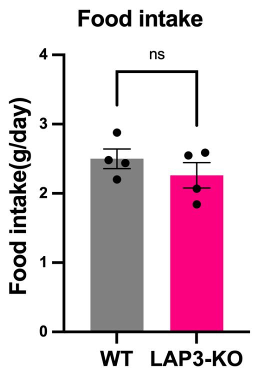


**Supplementary Figure 2. Food intake is unchanged in female LAP3-KO mice**.

Daily food intake measured during indirect calorimetry in female WT (n = 4) and LAP3-KO (n = 4) mice. Data are presented as mean ± SEM. Statistical significance was assessed using an unpaired Student's t-test.

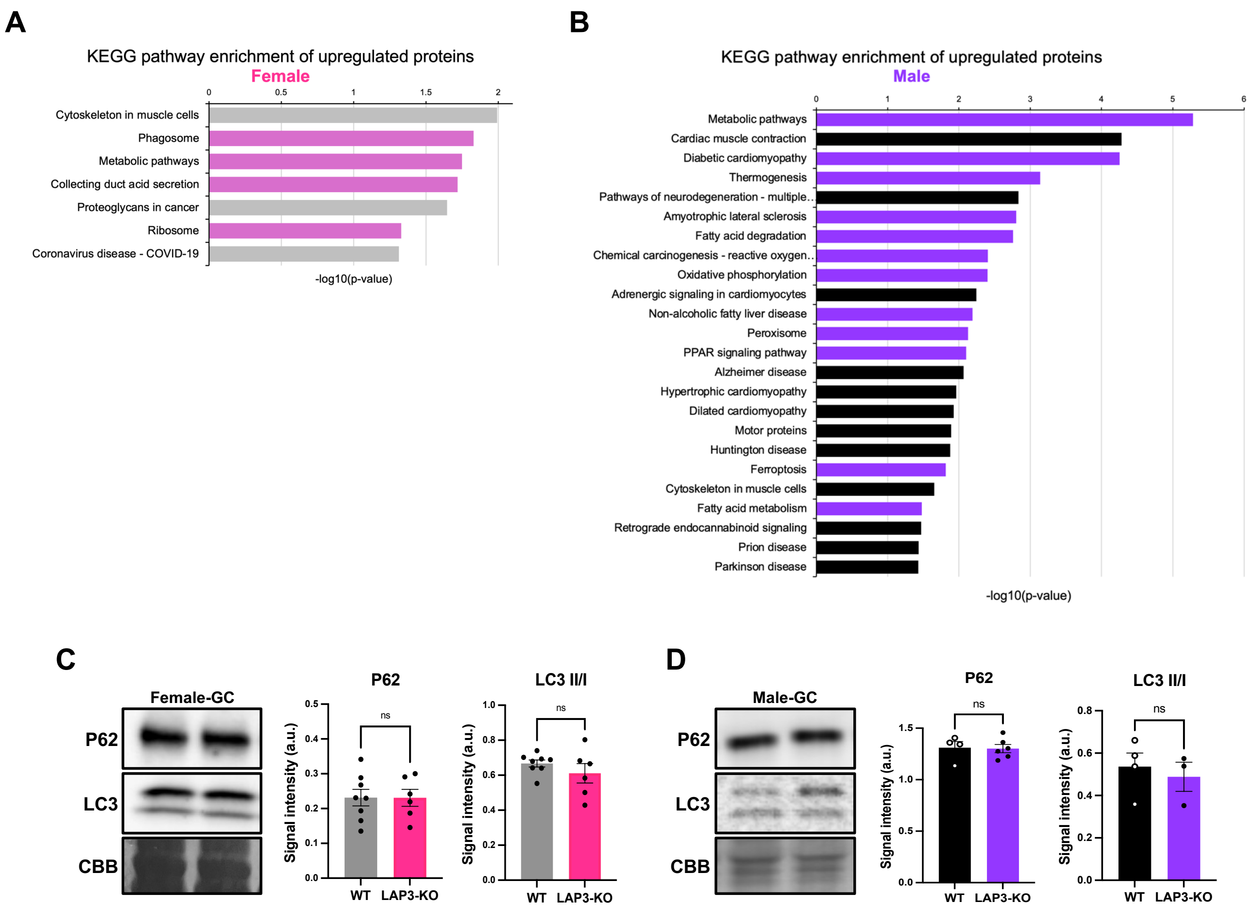


**Supplementary Figure 3. KEGG pathway analysis and autophagy-related protein expression in LAP3-KO skeletal muscle.** (A, B) KEGG pathway enrichment analysis of differentially expressed proteins in skeletal muscle from female (A) and male (B) LAP3-KO mice. (C, D) Representative western blots and quantification of p62 and LC3 expression in the gastrocnemius muscle of female (C) and male (D) WT and LAP3-KO mice. Data are presented as mean ± SEM (n = 3–8 mice per group). Statistical significance was assessed using an unpaired Student's t-test. $P < 0.05$.

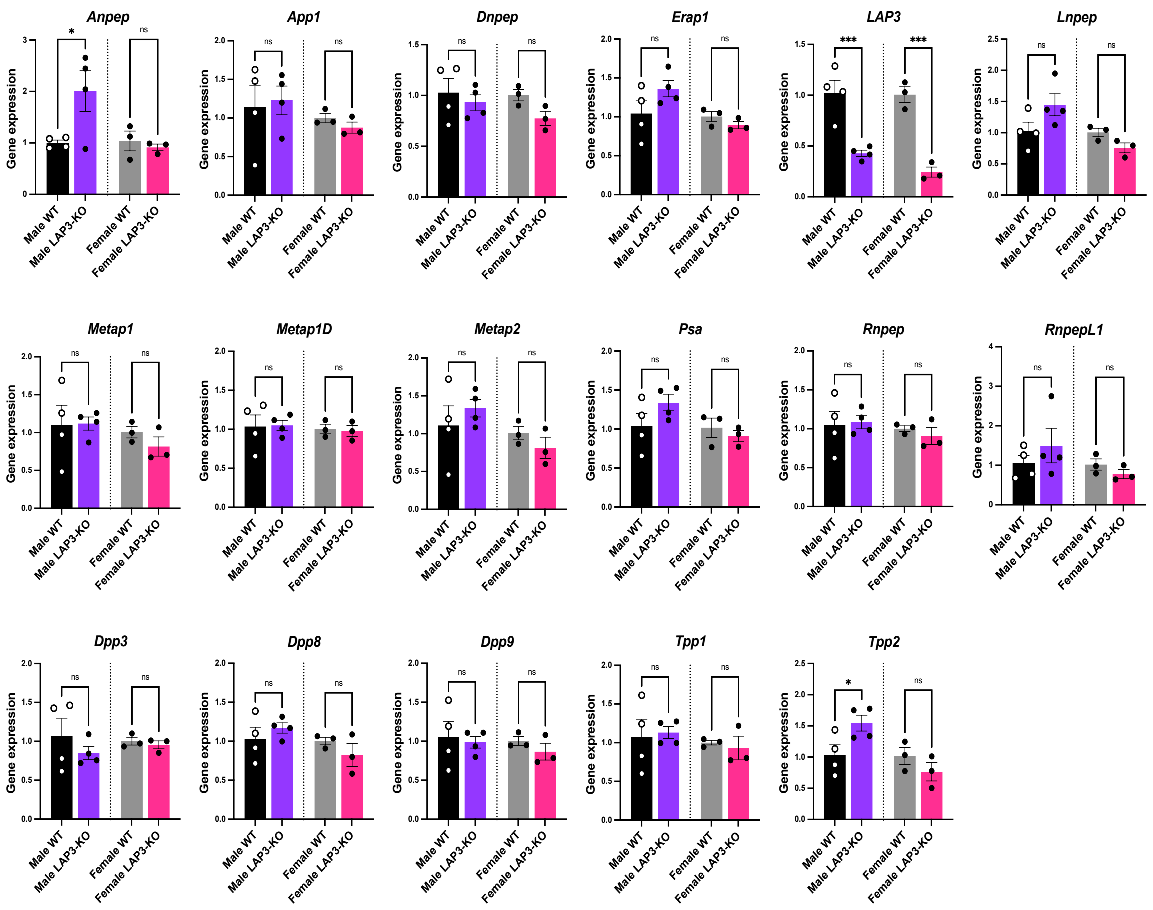


**Supplementary Figure 4. Expression analysis of aminopeptidase-related genes in the gastrocnemius muscle.** Quantitative PCR analysis of aminopeptidase-related genes in the gastrocnemius muscle of female and male WT and LAP3-KO mice. Data are presented as mean ± SEM (n = 3–4 mice per group). Statistical significance was assessed using an unpaired Student's t-test. *$P < 0.05$, ***$P < 0.001$.
